# Molecular Coevolutionary Drive

**DOI:** 10.1101/2023.04.22.537930

**Authors:** Michael Lynch

## Abstract

Most aspects of the molecular biology of cells involve tightly coordinated intermolecular interactions requiring specific recognition at the nucleotide and/or amino-acid levels. This has led to long-standing interest in the degree to which constraints on interacting molecules result in conserved vs. accelerated rates of sequence evolution, with arguments commonly being made that molecular coevolution can proceed at rates exceeding the neutral expectation. Here, a fairly general model is introduced to evaluate the degree to which the rate of evolution at functionally interacting sites is influenced by effective population sizes (*N_e_*), mutation rates, strength of selection, and the magnitude of recombination between sites. This theory is of particular relevance to matters associated with interactions between organelle- and nuclear-encoded proteins, as the two genomic environments often exhibit dramatic differences in the power of mutation and drift. Although genes within low *N_e_* environments can drive the rate of evolution of partner genes experiencing higher *N_e_*, rates exceeding the neutral expectation require that the former also have an elevated mutation rate. Testable predictions, some counterintuitive, are presented on how patterns of coevolutionary rates should depend on the relative intensities of drift, selection, and mutation.

**Significance Statement:** A wide variety of features at the cellular level involve precise interactions between participating molecular partners, thereby requiring coordinated co-evolutionary changes within lineages. The same is true for ecological interactions between species. Although there has been much speculation on how such constraints might drive molecular evolutionary rates beyond the neutral expectation, there has been little formal evolutionary theory to evaluate the generality of such claims. Here, a general framework is developed for ascertaining how rates of sequence evolution depend on the population-genetic environments of both interacting partner molecules. Although the features of one site can indeed drive the evolution of the other, only under restrictive conditions does this process push rates beyond the neutral expectation.

---

Molecular coevolution across nucleotide sites within species is pervasive. For example, the vast majority of proteins assemble as multimers, with homomers requiring the coordinated evolution of specific residues on binding interfaces within single proteins, and heteromers requiring coordination across genetic loci. For proper gene expression, key residues of transcription factors must closely match specific binding motifs on DNA, and for accurate translation, tRNA amino-acyl synthetases must specifically bind cognate tRNAs loaded with the appropriate amino acid. Signal-transduction systems relay specific messages between receptors and response regulators via precise binding interactions, and vesicle trafficking in eukaryotes involves multiple layers of protein-protein interactions to achieve delivery of specific cargoes to appropriate locations. Proper protein folding often requires accurate recognition of client molecules by chaperones. Bacterial plasmids often ensure their own survival by harboring precisely interacting toxin-antitoxin systems.

These few examples suffice to illustrate that much of cell biology is based on intragenomic molecular coevolution, just as much of community ecology is driven by interspecies co-evolution. A key difference with intragenomic coevolution is that the participating partners are bound together through shared inheritance, genetic linkage and/or common population-genetic environments. What remains unclear, however, is the degree to which such features stabilize vs. drive molecular evolution. On the one hand, one might expect coevolution to slow down rates of evolution, as the nucleotide content of an individual site constrains the domain of acceptable changes at other interacting sites. On the other hand, such interactions might induce an acceleration of coordinated changes, as when a mildly deleterious change at one site enhances the strength of selection for a compensatory change at another site. There is a long history of interest in this problem, ranging from the covarion hypothesis of Fitch and Markowitz (1970) to the Stokes-shift hypothesis of Pollock et al. (2012).

Some have argued that certain kinds of cellular evolution result in drive-like processes that encourage rates of molecular evolution even beyond the neutral expectation. For example, the centromere-drive hypothesis postulates that, in species with “female meiosis,” wherein only one of four meiotic products is delivered to an egg, centromere features that facilitate chromosome delivery into gametes will be strongly promoted through females, with negative side effects on male meiosis driving the emergence of secondary mutations to suppress the drive process (Henikoff et al. 2001; Malik and Henikoff 2001). Similar ideas are found throughout the wider literature on meiotic drive (Lindholm et al. 2016; Bracewell et al. 2019). Considerable attention has also been given to the idea that owing to elevated rates of mutation and a higher vulnerability to genetic drift, the accumulation of mildly deleterious mutations in organelle genes can impose selection for compensatory changes in the products of interacting nuclear-encoded genes (Rand et al. 2004; Sloan et al. 2018). In addition, coevolutionary drive has been invoked as an explanation for the rapid and coordinated evolution of proteins involved in mate recognition, e.g., sperm recognition and entry (Clark et al. 2009), and mating pheromones and their receptors (Luporini et al. 2005; Tsuchikane et al. 2005; Martin et al. 2011).

What remains unclear is whether the expectations of these largely verbal arguments hold up to more formal scrutiny when the underlying population-genetic processes are taken into consideration. Here, simple models are introduced to evaluate the conditions under which coevolution is expected to drive molecular evolution to rates approaching or exceeding the neutral expectation. In the absence of selection, the rate of molecular evolution per nucleotide site is equal to the mutation rate *u* (Kimura 1983), whereas a predominance of purifying selection to remove mutations or of positive selection for change leads to rates of fixation smaller than or larger than *u*, respectively. This general principle underlies the widespread empirical use of *d_N_ /d_S_*, where *d_N_* is the rate of evolution at nonsynonymous (amino-acid altering) nucleotide sites and *d_S_* is the rate of evolution at synonymous (and putatively neutral) sites, both of which are readily estimated by comparing gene-sequence data from closely related species. Thus, the following analyses focus on expected rates of evolution at key functional sites scaled to the mutation rate.

## RESULTS

### General two-locus model

Molecular coevolution involves epistasis, as the genotypic fitnesses at each locus depend on the genetic constitution at the other. The matter is evaluated here with a simple model involving two nucleotide sites (or genetic loci), each with two alternative states, **a/A** and **b/B** respectively. A haploid system is assumed (extension to diploidy only requires that population sizes be multiplied by 2, provided within-locus effects are additive), and the fitnesses for the four haplotypes **ab**, **aB**, **Ab**, and **AB** are taken to be 1, 1 *− s*_1_, 1 *− s*_2_, and 1 *− s*_4_ (Figure 1). Thus, assuming *s*_4_ = 0, a **b** *→* **B** mutation would restore the optimal state of fitness when occurring on a **Ab** background, and similarly for an **a** *→* **A** mutation on an **aB** background. A simple example of such a model involves stem bonds in RNA molecules, where A:T and G:C are viewed as alternative high-fitness states, with G:T and A:C being the only viable low-fitness intermediates. However, in its most general form, the model can be used to explore the molecular coevolutionary features of different interacting molecules, including those encoded in different genomic environments (e.g., organelle vs. nuclear). In addition with a change in sign of *s*, any haplotype can have a fitness exceeding that of the normalized **ab** type (here a positive *s* designates a deleterious mutation).

**Figure 1.**
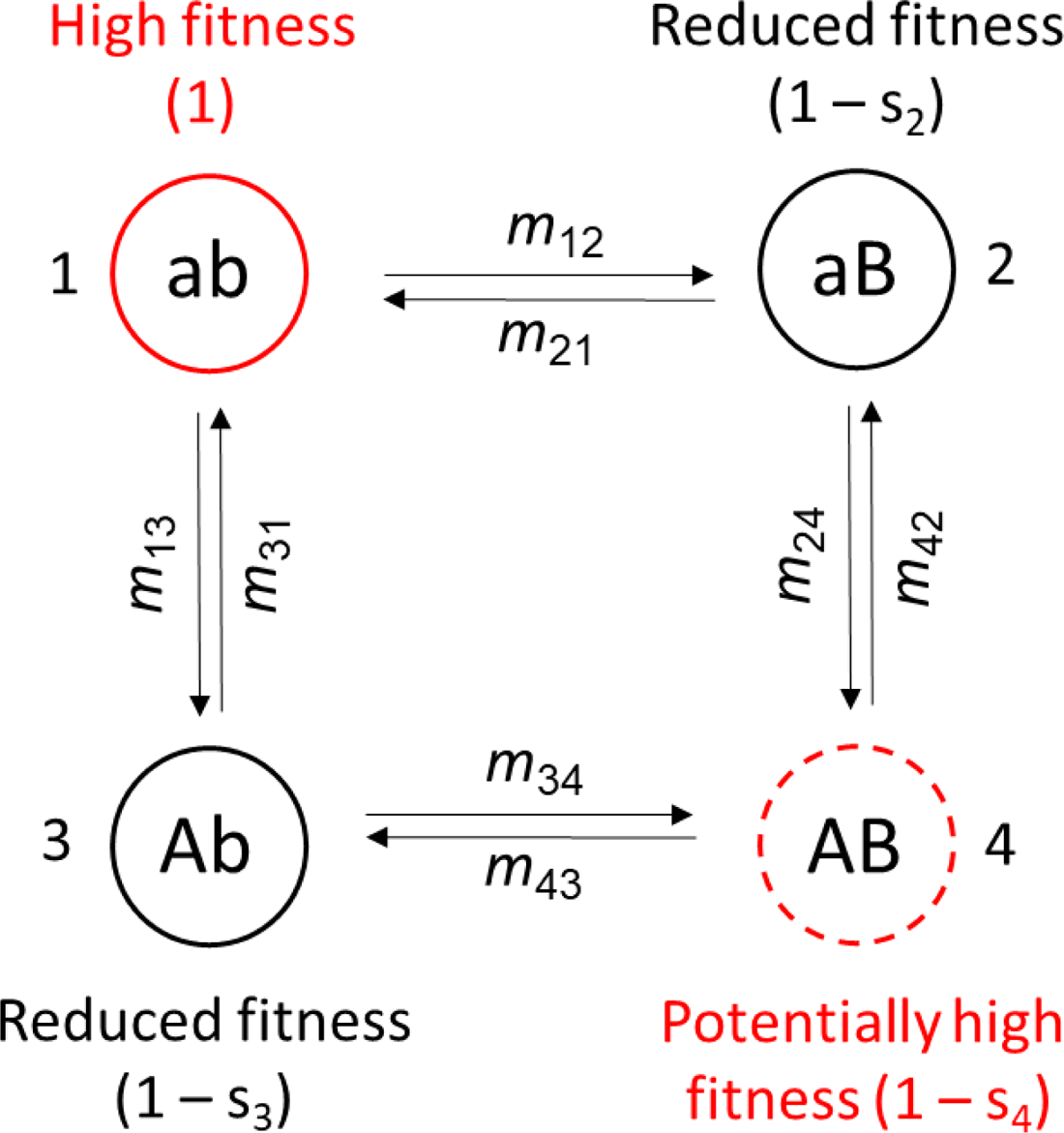
Flow diagram for four haplotype states, with the *m_xy_* denoting population-level rates of transition from one state of fixation to another. The four states (*x,y* = 1 …, 4) are denoted next to therr genotypic designa ions. The ab haplotype is arbitrarily designated as the one with the highest fitness (scaled to equal 1) but the modd is otherwise fully general, i.e., there is no e sential ordering of magnitudes of *s_2_ s_3_, and s_4_*. In the case of pmely compensatory mutations the AB haplotype has fitness equivalent to hat of ab 1.e., s_4_ = 0), and in the case of symmetry *s* = *s_2_ = s_3_*.

Aspects of this model were studied by Kimura (1985), but over more limited parameter space than explored here, and with three restrictive assumptions: 1) that both sites have identical mutation and drift properties; 2) that the strength of selection exceeds that of mutation; and 3) that mutations are irreversible. Extensions were made by Higgs (1998) to allow for reversible mutations, but again under the assumptions of constant population sizes and mutation rates. Both studies reduced a double-diffusion process to a single dimension to estimate the time to transition from one fixed beneficial state to the other. Below, more general expressions are given for the average rates of nucleotide substitution at both sites, allowing exploration of the full range of population-genetic environments and revealing behavior not previously appreciated.

The goal here is to evaluate the long-term rate of sequence evolution for interacting sites. To accomplish this, reversible mutations are allowed for, such that over time, the population will wander among alternative evolutionary states dominated by one of the four haplotypes. The residence time in each category and the transition rates between states depend on the fitnesses of the intermediate types, as well as on the effective population sizes and mutation rates associated with each site. From a knowledge of this quasi-steady-state distribution of alternative states and the forms of the transition coefficients, one can then derive the long-term average substitution rates at the two sites. Although only two biallelic sites are pursued in this initial analysis, the results should also apply to an entire system of interacting pairs, provided that each pair of interacting sites evolves independently of other such pairs.

We start with the assumption that fixations of new mutations occur sequentially, such that the population almost always resides in one of the four monomorphic states, with transitions only occurring between adjacent haplotype states in a stepwise fashion. Such a system will then evolve to a steady-state distribution of alternative states, defined by the relative magnitudes of the per-generation transition coefficients denoted in Figure 1. The probabilities of a population residing in the four alternative states over a long evolutionary time period are:

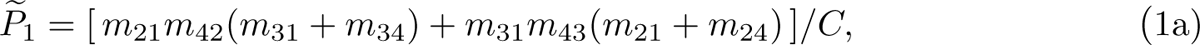

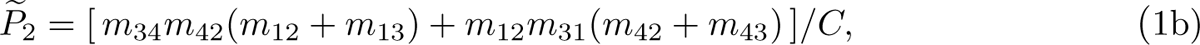

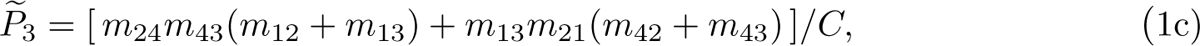

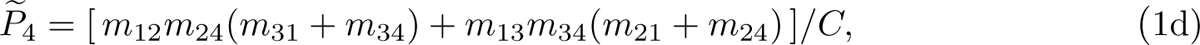

where *C* is a normalization constant equal to the sum of the numerators in Equations 1a-d. Although the algebra is tedious, there is a simple heuristic way of obtaining these results. The numerator for each probability is equal to the sum of four terms, each a product of the three transition coefficients jointly pointing towards the haplotype state (two chains of three consecutive coefficients, and two involving one transition pointing from one direction and a chain of two pointing from the other direction). This rule of thumb may be useful for obtaining steady-state frequencies involving more than two states (e.g., the eight haplotypes with a three-site system), although the number of contributing pathways increases rapidly with the number of sites.

Each transition coefficient is equal to the product of the rate of origin and probability of fixation of the mutation type. Denoting mutation rates and population sizes for the two loci as *u_A_* and *u_B_*, and *N_A_* and *N_B_*, respectively, the population-level rates of origin of mutations for the two loci are *U_A_*= *N_A_u_A_* and *U_B_* = *N_B_u_B_*. The probability of fixation of a mutation arising at locus *x* is given by

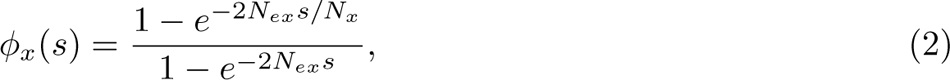

(Maĺecot 1952; Kimura 1957), where *N_ex_* and *N_x_* denote the effective and absolute population sizes for locus *x*, and the value of *s* depends on the difference in fitness between the mutant haplotype and the originating type (as outlined in Figure 1). For example, *m*_12_, which denotes the fixation of a deleterious mutation arising at the second locus on a **ab** background, is equal to *U_B_ · φ_B_*(*−s*_2_). In contrast, *m*_24_ denotes a mutation arising at the first locus (on a **aB** background) and is equal to *U_A_ · φ_A_*(*s*_2_ *− s*_4_).

Substantial simplification of Equations 1a-d is possible in a number of cases. Consider, for example, the situation of complete symmetry, with the two intermediate haplotypes (**aB** and **Ab**) having equivalent reductions in fitness (*s* = *s*_2_ = *s*_3_), and the two end types (**ab** and **AB**) having equivalent higher fitness (*s*_4_ = 0). Depending on whether the transitions are beneficial (*b*) or deleterious (*d*), each new mutation has a probability of fixation of

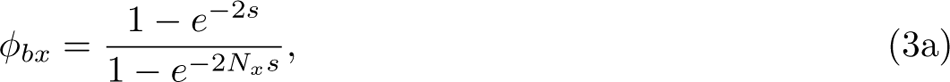

Or

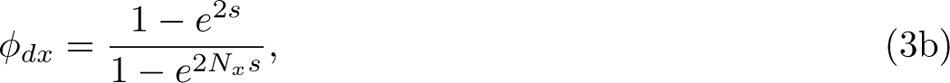

where *x* denotes the locus of mutational origin. Here, for purposes of illustration, it is assumed that effective and absolute population sizes are equal to each other within sites but not necessarily between the two loci. If this is not the case, the appropriate modifications (as in Equation 2) need to be implemented; in effect, the necessary changes are simply equivalent to substituting for *s* an effective selection coefficient equal to *N_e_s/N*. For this special case of symmetrical fitnesses, the system of Equations 1a-b reduces to the following long-term steady-state probabilities of the population residing in the four alternative states:

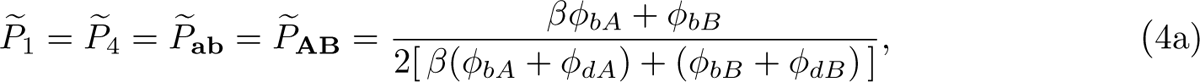

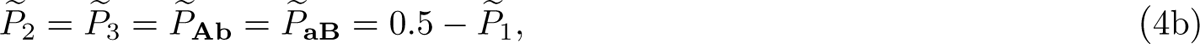

where *β* = *U_A_/U_B_*.

General expressions for the mean rates of substitution per site relative to their neutral expectations are obtained by weighting the rates of transition between alternative states by the originating haplotype probabilities, and dividing by site-specific mutation rates,

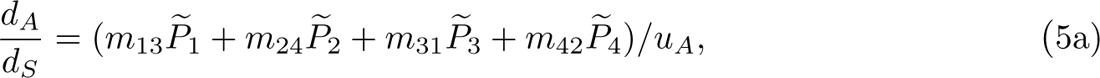

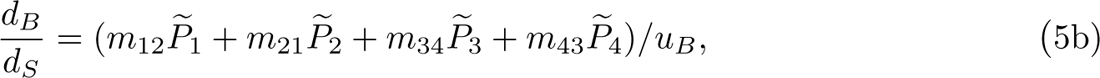

which for the case of symmetrical fitnesses reduce to

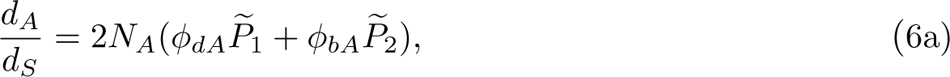

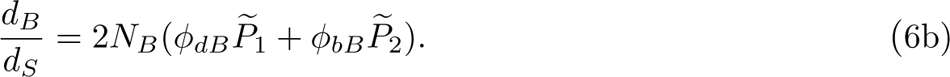

To evaluate the adequacy of these sequential-model approximations, as well as to obtain more general results, computer simulations were run over the full range of mutation and recombination rates, population sizes, and selection intensities known across the Tree of Life (Lynch 2007; Lynch et al. 2016; Lynch and Trickovic 2020). These followed a standard Wright-Fisher structure with consecutive episodes of deterministic mutation (at rates *u_A_*and *u_B_*), recombination (at rate *c* between sites), and selection (using the coefficients *s*_2_, *s*_3_, and *s*_4_), followed by multinomial sampling of the four haplotype frequencies (to incorporate random genetic drift, with population sizes ranging from *N* = 10^3^ to 10^9^). To maintain reasonable computational times at large *N*, some down-scaling of *N* was done, while keeping *Nu*, *Ns,* and *Nc* constant, which retains the proper scaling of stochastic relative to deterministic processes. After verifying that such rescaling did not influence the equilibrium results, simulations proceeded for 10^7^*N* to 10^9^*N* generations for each parameter set, with population status being monitored every *N/*10 generations. The final compilation of results for each set of population-genetic conditions yielded information on the long-term mean frequencies of the four haplotypes, *P_x_,* along with the evolutionary rates (summed frequencies of bidirectional transitions between alternative alleles) for each site. For practical reasons, fixations were inferred to have occurred when a haplotype with frequency *>* 0.999 gave rise to another haplotype above the same threshold.

### Complete linkage

We start with the situation in which the two loci are completely linked, as will often be true for nucleotide sites within the same gene or for any pair of sites on a nonrecombining chromosome (commonly believed to be the case within organelle genomes, and always the case with an asexual). Under these conditions, *N* = *N_A_*= *N_B_,* and *u* = *u_A_*= *u_B_.* In the case of complete symmetry, *φ_b_*= *φ_bA_* = *φ_bB_, φ_d_* = *φ_dA_*= *φ_dB_*, and Equations 6a,b for the sequential model reduce to

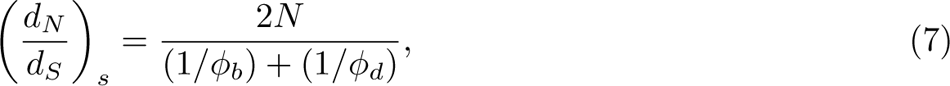

for both loci. This expression, which is equivalent to *N* times the harmonic mean of the beneficial and deleterious fixation probabilities for new mutations, is independent of the mutation rate, and further reduces to 4*Ns · e^−^*^2^*^Ns^* for *Ns >* 1. There is then no situation in which the rate of evolution exceeds the neutral expectation, as *d_N_ /d_S_ →* 1 for small *Ns* and *→* 0 for large *Ns*, with the shift in behavior occurring at *Ns ∼* 1.

As population sizes become large, the assumptions of the sequential model break down because deleterious haplotypes maintained at low frequency by selection-mutation balance can acquire secondary (compensatory) mutations. This enables simultaneous fixations at both loci in a neutral fashion, e.g., an **AB** haplotype arising by two mutations in a population dominated by **ab** haplotypes. Such a process is commonly referred to as stochastic tunneling (Komarova et al. 2003; Iwasa et al. 2004; Weissman et al. 2009; Lynch and Abegg 2010). As an example, for the case of symmetrical fitnesses (*s*_2_ = *s*_3_ = *s,* and *s*_4_ = 0), approximately half of the time, the population will be dominated by the **ab** haplotype, with deleterious **Ab** and **aB** haplotypes each being maintained at frequencies near *u/*(2*u* + *s*) by selection-mutation balance (*∼ u/s* for *s » u*). Multiplying these frequencies by the number of individuals, noting that compensatory mutations (at the second locus) arise and fix at rates *u* and 1*/N*, and taking the products of terms and summing through both pathways yields a rate of transition from dominance by the **ab** to the **AB** haplotype equal to 2*u*^2^*/*(2*u* + *s*). The same rate applies in the opposite direction, and dividing by the mutation rate yields the scaled tunneling rate

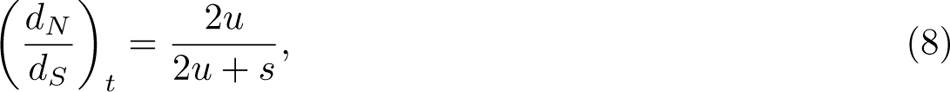

in the limit of large *N* (*∼* 2*u/s* for *s » u*). Unlike the situation in the sequential-fixation regime, the scaled tunneling rate depends on the mutation rate, but is independent of the population size.

Taking into consideration both paths to fixation, and noting that as a first-order approximation, the stochastic-tunneling regime requires that the power of selection exceed that of drift, a composite estimate of the total substitution rate per locus (relative to the neutral expectation) is

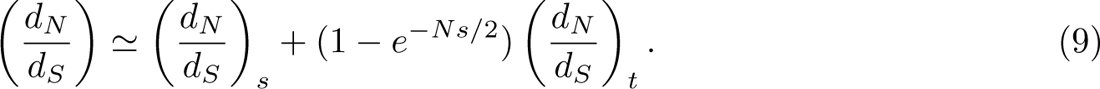

For low mutation rates, *u* = 10*^−^*^9^, which are in accord with per-site estimates for a wide range of microbes (Lynch et al. 2016), the predictions from Equation 9 are in close agreement with observations derived from computer simulations (Figure 2). At *u* = 10*^−^*^7^, above the known upper limit for cellular organisms, the simulation estimates fall below Equation 9 when *s/u ≤* 1 and *Nu >* 0.1. However, such deviations are simply a consequence of the failure to always properly identify true genealogical fixation events with haplotype frequency data when high levels of polymorphism are maintained by mutation pressure.

**Figure 2.**
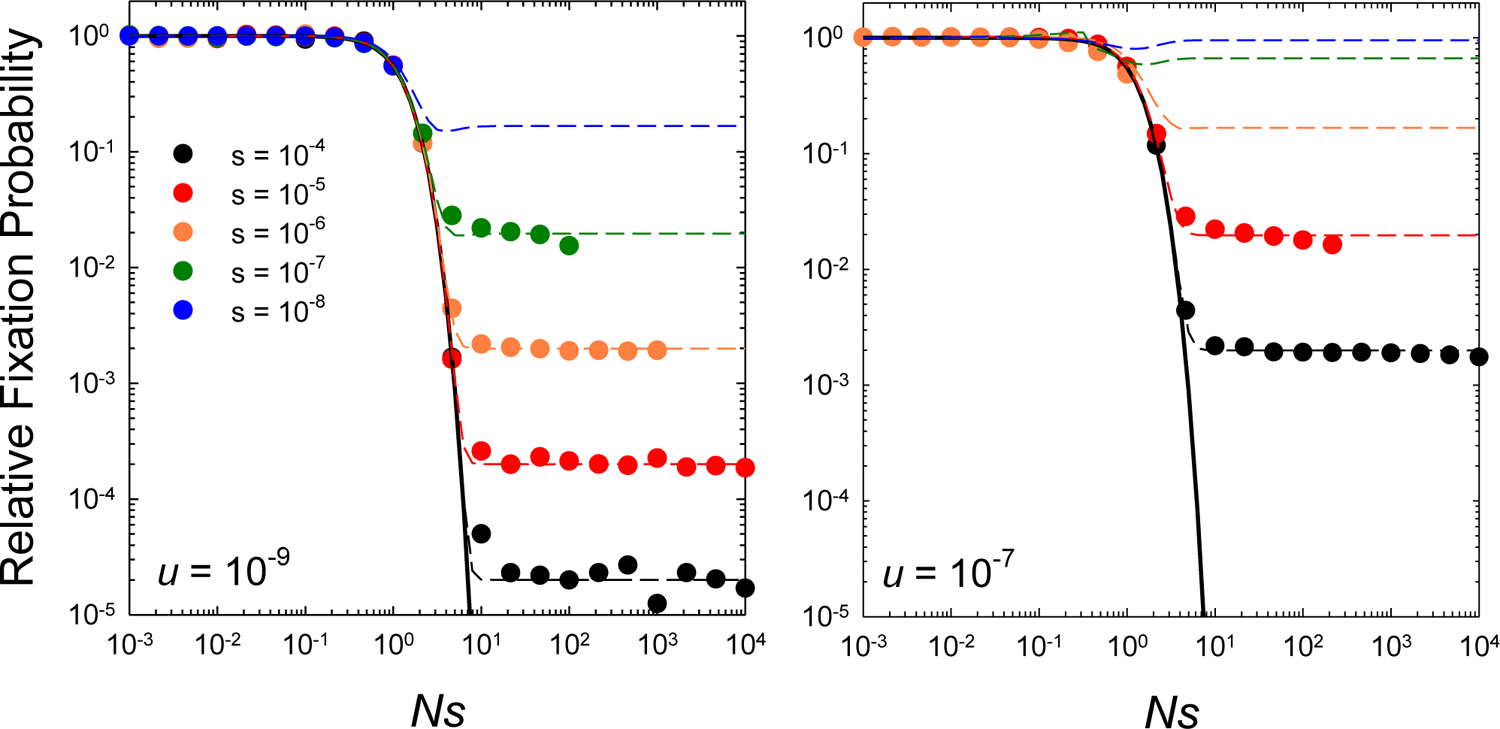
Rates of substitution at the two sites scaled to the neutral expectation as a function of *N*s (the ratio of the strength of selection to drift), for the case of complete linkage with both sites experiencing equal mutation rates, selection coefficients (*s* = *s_2_* = *s*_3,_ and *s*_4_ = 0), and population sizes. The data points denote computer-simulation results, obtained over a range of *N*_e_ from 10^3^ to 10^9^ for each set of *s* and *u*. The solid black line denotes the sequential-model prediction, which is independent of the mutation rate (Equation 7), whereas the horizontal dashed lines extend to the stochastic-tunneling domain (Equation 9), and do depend on the mutation rate. Simulation data on fixation rates are only given when the population level mutation rate per site *Nu* < 0.1 and the deterministic expected frequency of the deleterious type ≌ *u/s* < 0.1, as otherwise recurrent mutations make it difficult to accurately ascertain fixation events from observed allele frequencies that become bounded away from 1.0.

The general conclusion to be drawn for completely linked sites is that in the region of 1 *< Ns <* 10 there is a precipitous shift in the behavior of *d_N_ /d_S_* from the expectations under the sequential model (*Ns <* 1) to those under the tunneling model (*Ns >* 10). As expected from Equation 8, the magnitude of this drop-off increases with larger *s/u* because intermediate-state haplotype frequencies are reduced when selection is stronger. As the intensity of selection declines below the mutation rate, the scaled tunneling rate 2*u/*(2*u* + *s*) *→* 1, and the population spends nearly equal time in the four alternative states.

Extension of these results to other fitness schemes are presented in the Supplemental Material. The key point is that, for the case of complete linkage, the main conclusions derived for the case of symmetrical fitnesses are retained. Regardless of the level of asymmetry in fitnesses, both sites evolve at identical relative rates and *d_N_ /d_S_ ≤* 1, i. e., molecular coevolution does not drive the rate of evolution beyond the neutral expectation. For the case in which the two intermediate states have different fitnesses (*s*_2_ */*= *s*_3_) but the end states have equal fitness (*s*_4_ = 0), the preceding conclusions continue to hold, except that the transition from the sequential to the tunneling domains extends over a longer range of *N*, owing to the fact that the two sites have different ranges of effective neutrality. For the case in which *s*_2_ = *s*_3_ but *s*_4_ *>* 0 (one end state has a higher fitness advantage than the other), the relative rate of evolution continuously declines with increasing *N*, owing to the fact that the population resides almost entirely in the beneficial end state, asymptotically diminishing the probability of tunneling towards zero.

### Free recombination

In sexually reproducing species, for nucleotide sites on different chromosome arms, including in organellevs. nuclear-encoded genes, segregation across sexual generations will be nearly completely independent, rendering the process of stochastic tunneling essentially inoperable (Lynch 2010; Weissman et al. 2010). Consider a large population predominately in a beneficial **AB** state, with the **Ab** and **aB** haplotypes maintained by selection-mutation balance. Although recombination can readily create an **ab** haplotype out of an **Ab/aB** pair, almost all haplotypes subsequently recombining with the newly arisen **ab** haplotype will be of type **AB**, rapidly converting it back to **Ab** and/or **aB** types. This barrier to evolutionary transitions is most apparent when mutation rates and effective population sizes are constant across sites, in which case the scaled substitution rate is closely approximated by Equation 7, again independent of the mutation rate and largely a function of *Ns* (Figure 3A).

**Figure 3.**
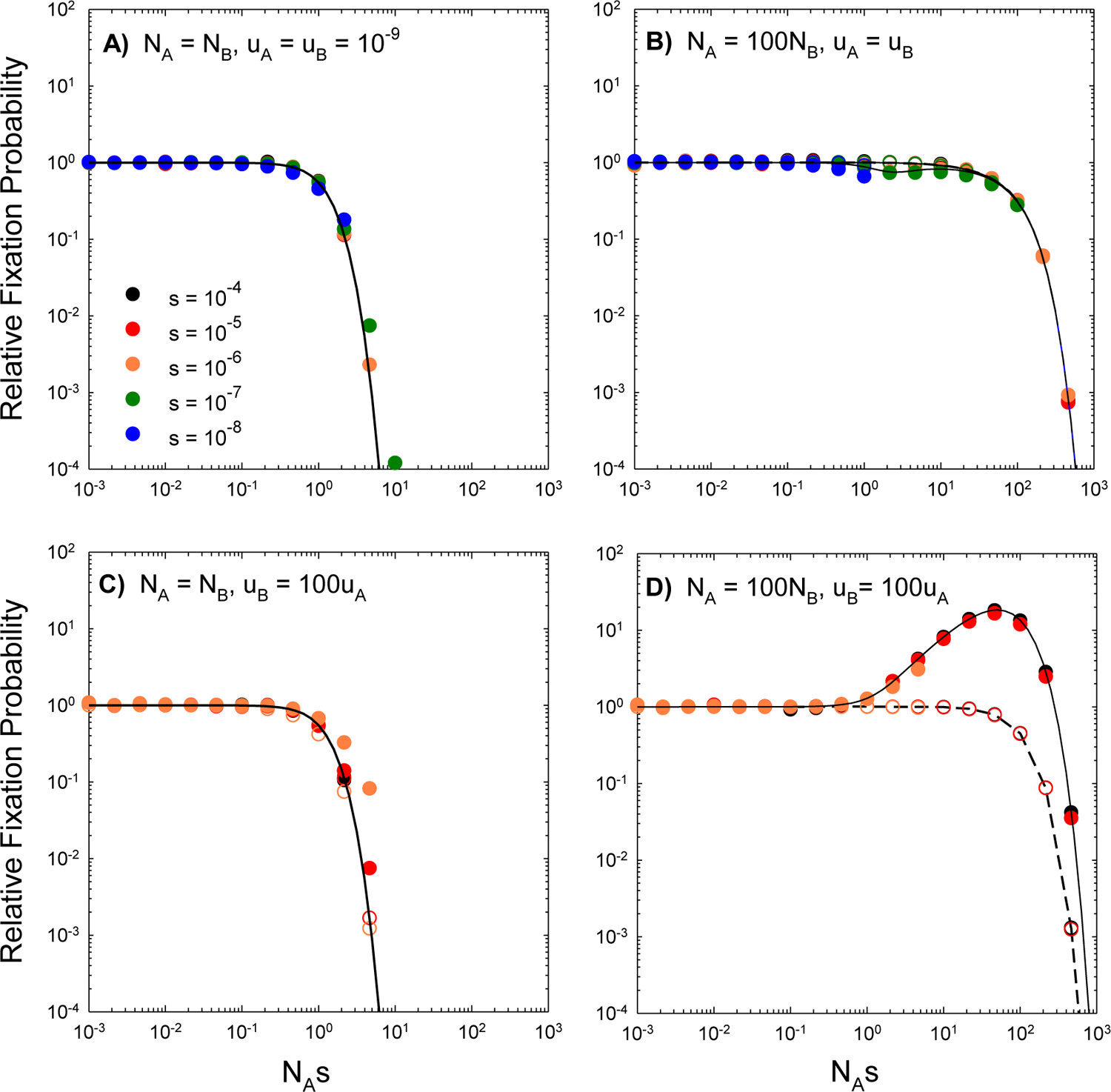
Scaled rates of substitution at the two sites relative to the neutral expectation as a function of *N*_A_s (the ratio of the strength of selection to drift at the A/a locus), for the case of free recombination, with various combinations of effective population sizes and mutation rates between sites. Solid and open points are simulation results for sites A and B, respectively. Continuous lines denote the theoretical expectations described in the text. In all cases, *uA* = 10^-9^, and for any set of conditions simulations were performed over a range of *N*_A_ from 10^3^ to 10^9^.

If population sizes and/or mutation rates differ among loci, as will often be the case when the two sites reside in nuclear vs. organelle environments, the behavior of the system is still well-described by Equations 5a,b. To appreciate the nuances that can arise, however, consider the simplest case of symmetry in which *s* = *s*_2_ = *s*_3_ and *s*_4_ = 0 and Equations 6a,b apply (Supplemental Text). When mutation rates are constant but population sizes differ, there are three domains of behavior (Figure 3B): 1) for *N_A_, N_B_ <* 1*/s*, the regime of complete effective neutrality, the scaled rates of evolution *∼* 1.0 for both loci; 2) for *N_A_ >* 1*/s* but *N_B_ <* 1*/s*, the first locus is under efficient selection and the second is effectively neutral, but the scaled rates for both loci remain *∼* 1.0; and 3) for *N_A_, N_B_ >* 1*/s*, such that both loci are under efficient selection,

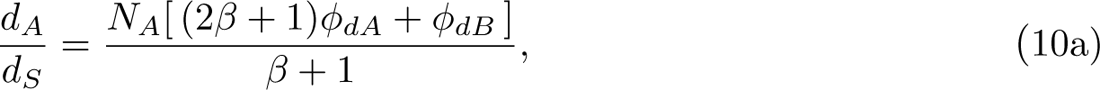

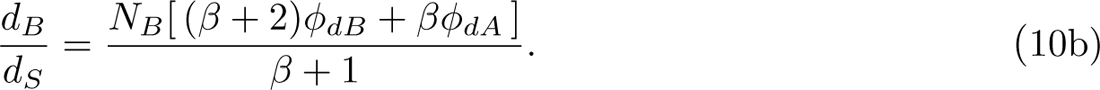

In the latter case, which applies even when mutation rates differ, both *d_A_/d_S_* and *d_B_/d_s_*are kept low by strong selection for beneficial alleles, with the dynamics being largely determined by rare fixations of deleterious alleles and the subsequent compensatory responses. Under these conditions, the smaller of the two effective population sizes simply extends the regime of apparent effective neutrality (*d_A_/d_S_* = *d_B_/d_S_ ∼* 1), with the scaled rates of evolution being nearly equal for both sites (Figure 3B). In effect, the elevated rate of fixation of deleterious alleles at the site with smaller *N* drives the scaled rate of evolution at the large-*N* site to a similar level by promoting the fixation of compensatory mutations.

When *N* is constant across sites but mutation rates differ, Equations 5a,b both reduce to Equation 7, showing that between-site variation in the mutation rate alone has no impact on the scaled rates of evolution in the case of free recombination (Figure 3C). In this case, one site produces more deleterious mutations than the other, but because the fixation probabilities are identical at both sites, both produce restorative mutations at rates proportional to their own mutation rates.

Qualitatively different behavior arises when both *N* and *u* vary between sites. Consider the situation in which one site experiences a lower effective population size but higher mutation rate, as seems to frequently occur with metazoan mitochondria (Lynch 2007). Under these conditions, the site with the lower *N* behaves in the same manner as discussed above, with the shoulder on its scaled rate function extending to the point appropriate for the smaller population size (Figure 3D). In contrast, the scaled evolutionary rate at the site with large *N* /small *u* is elevated above the neutral expectation in a large fraction of the domain in which *N_A_s >* 1 and *N_B_s <* 1, a point previously noticed by Osada and Akashi (2012).

To see why this occurs, consider the extreme case in which *φ_bA_ ∼* 2*s, φ_dA_ ∼* 0, and *φ_bB_* = *φ_dB_ ∼* 1*/N_B_*, i.e., highly efficient selection at locus A and effective neutrality at locus B. In this domain,

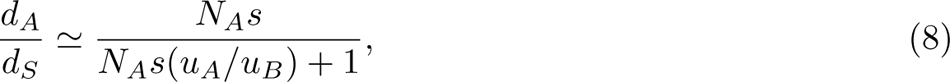

which for *N_A_s*(*u_A_/u_B_*) *«* 1 is approximately *N_A_s*, as can be seen in Figure 3D. This approximation can be further understood by considering that under the conditions noted, the low-*N_e_* site spends half the time in the **B** vs. **b** state, with the high-*N_e_*locus mutating at rate *N_A_u_A_* and fixing with probability 2*s*; dividing the product of these three terms by *u_A_*returns *d_A_/d_S_∼ N_A_s.* (Note that under the opposite mutational condition (*u_A_» u_B_*), *d_A_/d_S_ ∼ u_B_/u_A_* because evolution at low-*N_e_* site proceeds at its neutral rate *u_B_*, eliciting the same rate of change at the high-*N_e_*site, as compensatory mutations arise at a higher rate than reversions).

Allowing for asymmetry in the strength of selection does not greatly alter the behavior of recombining systems (Figure 4). First, when mutation rates are equal, but there is asymmetry in population sizes (Figure 4A), the site with small *N* again drives the process and always has the higher scaled evolutionary rate, as it is most susceptible to the fixation of mildly deleterious mutations. However,the site with higher *N* (in this case A) exhibits a nonmonotonic scaling of the scaled fixation probability with population size, initially declining with increasing *N_A_*, then increasing, and then finally monotonically decreasing.

**Figure 4.**
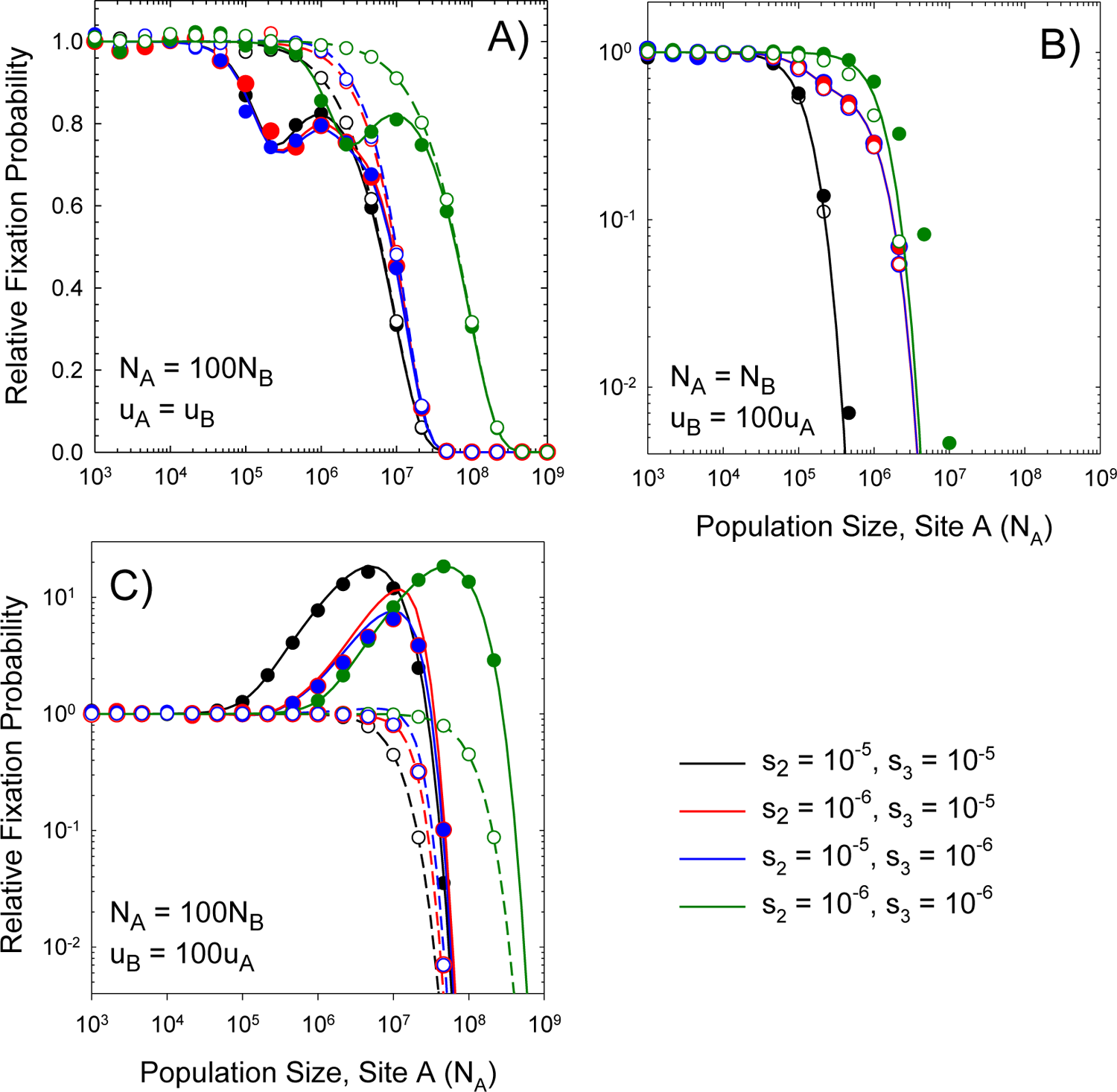
Scaled rates of substitution at the two sites relative to the neutral expectation as a function of *N_A_* (the population size at the A/a locus), for the case of free recombination and asymmetry in the strength of selection (*s*_2_ ≠ *s*_3_), with various combinations of effective population sizes and mutation rates between sites. In all cases, *uA* = 10^-9^. Solid and dashed lines, derived from theory, denote results for the A and B loci, respectively.

This even occurs when *s*2 = *s*_3_. The initial valley in the scaled fixation probability for site A occurs at *N_A_ ∼* 2*/s*_3_. To the left of this inflection point, the strength of selection relative to drift is weak but non-negligible at site A, which is incapable of fully responding to the challenges resulting from deleterious fixations at site B. Immediately to the right of the inflection point, site B is still evolving at the neutral rate, but selection at site A is efficient enough to nearly fully compensate for B-site fixations. Once both population sizes are sufficiently large, deleterious fixations are uniformly inhibited, and the scaled rates of evolution decline in an identical fashion at both sites.

Second, when population sizes are identical but mutation rates differ (Figure 4B), both sites have identical *d_N_ /d_S_,* which beings to decline monotonically with increasing *N* at a position determined by the strength of selection at both sites. Finally, when there are large *N* /small *u* and small *N* /large *u* sites, there is again a domain of intermediate population sizes in which the former has *d_N_ /d_S_ >* 1. Asymmetric selection simply alters the position of the peak.

### Intermediate levels of recombination

The preceding results provide accurate descriptions of the pace of sequence evolution in the extreme cases of zero and free recombination. To gain insight into how low the recombination rate needs to be for the complete-linkage results to hold, note that the mean time to fixation of a neutral mutation in a haploid population is 2*N_e_* generations and that it is reasonable to assume that linked sites have the same effective population sizes and mutation rates. Consider an **AB** haplotype, destined to fixation, arising by secondary mutation in a population currently dominated by **ab**. As the **AB** lineage proceeds to fixation, from start to finish, it will experience an average background **ab** frequency of 0.5, and if a recombination event involves an **ab/AB** pairing, 3*/*4 of the progeny will be non-**AB** type, so en route to fixation, the probability that a descendant of the original mutation will have avoided a recombination event by the time of fixation is on the order of *e^−^*^3^*^Nec/^*^4^. Although this argument ignores various aspects of the underlying dynamics of haplotype-frequency changes (including secondary restorative recombination events), the overall response of the scaled rate of evolution is closely approximated by multiplying the tunneling term in Equation 9 by *e^−Nec/^*^2^ (Supplemental Figure S2). Increasing *c* beyond 1*/*(2*N_e_*) eliminates the stochastic-tunneling domain, thereby imposing a strong barrier to evolution for *Ns >* 1.

Note that Kimura (1985) provided a complex expression for the role of recombination on the rate of stochastic tunneling, but his results suggesting only a moderate influence of recombination were based on the assumption that *c « s*, which will often not be the case, and hence are not strictly comparable with the results herein. Likewise, Higgs’ (1998) proposal of a recombination-rate barrier in the neighborhood of *c* = *u*^2^*/s* is inconsistent with the results here.

## DISCUSSION

The preceding results yield insight into several consequences of molecular coevolution between a pair of genomic sites. Nucleotide sites with mutually constrained fitness effects can be expected to undergo coevolutionary divergence among phylogenetic lineages to an extent that might ultimately give rise to interspecies incompatibilities (Coyne and Orr 2004). However, the degree to which intermolecular constraints enhance vs. curtail the rate of evolution depends greatly on the population-genetic environment, sometimes in counterintuitive ways. Even with constant selection pressures (*s*), patterns of evolutionary rates driven by molecular coevolutionary pressures can vary substantially, depending on variation in *N_e_*, *u*, and *c* between interacting partners, usually as ratios of pairs of evolutionary pressures, i.e., *N_e_s*, *N_e_c*, and *u/s*, at both loci.

Because interacting sites may comprise only a small to moderate fraction of the full sequences of interacting genes, the behavior of scaled evolutionary rates (*d_N_ /d_S_*) at the gene-wide level need not closely reflect the subset of sites experiencing compensatory mutation, although with adequate phylogenetic data spatial analyses across gene bodies should be capable of revealing the kinds of patterns predicted above. For the remainder of the discussion, the population size will be referred to as *N*, recalling that *N_e_ < N* has the effect of reducing the effective strength of selection to *N_e_s/N*.

For the case of complete linkage, both sites are expected to have identical effective sizes and mutation rates, and the predicted consequences of coevolution are quite general.

For symmetrical selection (*s*_2_ = *s*_3_ = *s*) and *Ns <* 1 (the domain of effective neutrality), *d_N_ /d_S_ ∼* 1, whereas for 1 *< Ns <* 5*, d_N_ /d_S_* is a decreasing function of *Ns* independent of the mutation rate (*u*). For *Ns >* 5 (the domain of stochastic tunneling), *d_N_ /d_S_* takes on a value defined by *u* and *s*, and likely continues to decline with increasing *N* owing to the negative association between population size and evolved mutation rates (Lynch et al. 2016).

Thus, for tightly linked sites (within closely spaced segments of genes, or anywhere within nonrecombining genomes), it is expected that: 1) scaled rates of evolution will decline with increasing *N* in different lineages experiencing identical selective pressures; 2) both participants will have matching evolutionary rates within phylogenetic lineages; and 3) *d_N_ /d_S_* will not exceed 1.0, i.e., there is no molecular coevolutionary drive beyond the neutral expectation. These conclusions hold even when there is an asymmetry in the strength of selection operating on sites, except that the transition between the two domains of behavior noted above occurs near the point where *N* is equal to the inverse of the smallest selection coefficient. That is, relaxed selection on one site induces a broader population-size domain in which both sites evolve in an effectively neutral fashion (as the more weakly selected site induces the need for compensatory evolution).

Free recombination alters these expectations for situations in which sites differ in effective population sizes and/or mutation rates (Lynch 2007). When differences in the power of drift exist between sites, as will commonly be the case for organelle vs. nuclear genomes, the site experiencing the smaller *N* dictates the rate of evolution at both sites, unless *N_A_, N_B_ »* 1*/s,* in which case molecular coevolution grinds to a halt, freezing the two sites into one particular configuration. Moreover, provided only *N* or only *u*, but not both, differ between sites, *d_N_ /d_S_* behaves in an essentially identical manner at both sites, as observed with linked sites. In effect, a sufficiently small-*N* site can evolve in a nearly neutral fashion (*d_N_ /d_S_ ∼* 1), with the overall system retaining high fitness as the associated large-*N* site is driven to fix compensatory mutations by selection. As a consequence, the latter has *d_N_ /d_S_ ∼* 1 well out into the range where its *Ns >* 1, a condition that is typically expected to lead to purifying selection.

With sufficiently high recombination rates, an asymmetry in the scaled patterns of evolution arises when both *N* and *u* differ between sites. Intermediate states of low fitness (**Ab** and **aB**) can be restored to high fitness by either a reversion at the prior site of mutation or by compensatory mutation at the alternative site. When one site has low *N* / high *u* (say site B) and the other high *N* / low *u* (site A), provided *s* is sufficiently low, site B evolves at the neutral rate by mutation pressure, driving the scaled rate of functional evolution at site A to levels as high as *N_A_s >* 1. This leads to the appearance of very strong positive selection on the site experiencing more efficient selection, although the process is completely driven by internal pressures. In the opposite situation, where there is high *N* / high *u* (site A) and low *N* / low *u* (site B) coupling, with the former being under efficient selection and the latter effectively neutral, *d_A_/d_S_∼ u_B_/u_A_ <* 1, the ratio of mutation rates at the two loci, rendering the appearance of stronger purifying selection on the high-*N* site. Thus, a simple change in the ratio of mutation rates in a coevolving system can lead to qualitatively different conclusions using *d_N_ /d_S_* as an indicator of the direction of selection (even when the magnitude of selection *s* is kept constant).

These results are relevant to the interpretation observations on DNA sequence evolution at putatively coevolving sites. For example, most methods for inferring coevolution from comparative sequence data rely on patterns of covariation of rates of base substitutions for pairs of sites/genes among different phylogenetic branches (e.g., de Juan et al. 2013; Cong et al. 2019). It has been argued that such covariation of *d_N_ /d_S_* reflects lineage-specific changes in selection pressures with parallel effects on members within specific cellular pathways (Clark and Aquadro 2010; Clark et al. 2013). However, although the use of *d_N_ /d_S_* sometimes controls for interspecies variation in mutation rates, this is not always the case with coevolving sites (as illustrated in several of the preceding formulations), and it does not remove the effects of population-size differences, which influence rates of functional-site evolution through their effects on fixation probabilities. Consider, for example, the situation for fully linked sites with constant *s* and *u* (Figure 2). Here, for populations straddling *Ns ∼* 5, both members of an interacting pair can jointly have relatively high or low *d_N_ /d_S_*, despite experiencing the same absolute strength of selection (*s*). Thus, phylogenetic correlations of *d_N_ /d_S_* between pairs of target sites/genes do not necessarily reflect temporal changes in absolute selection intensities.

Eukaryotic molecular complexes that derive their components from organelle- and nuclear-encoded genes have served as popular targets in studies of molecular coevolution, motivated by the substantial differences in *N* and *u* that can exist between these two genomes within species (Lynch 2007). In principle, some aspects of organelle bottlenecking and uniparental inheritance may also alter the efficiency of selection (Christie et al. 2017; Radzvilavicius et al. 2017; Edwards et al. 2021). Specific attention has been given to the idea that elevated rates of mutation and/or reduced effective population sizes in organelle genomes leads to the accumulation of mildly deleterious mutations, creating pressure for the fixation of compensatory mutations (Rand et al. 2004). Support for this idea derives from studies on the base-pairs in the stems of tRNAs and rRNAs in organelle vs. nuclear genomes (Lynch 1996, 1997; Oliveira et al. 2008; Meer et al. 2010; James et al. 2016). For such genic regions, the scaled rates of molecular evolution for nuclear genes are often two- to three-fold lower than for organelle genes, presumably because of reduced *N* experienced by the latter, and in accordance with the expectations for linked sites (Figure 2).

More closely connected to the theory for unlinked sites are molecular complexes consisting of nuclear- and mitochondrial-encoded proteins. Several studies have shown that phylogenetic lineages with rapidly evolving mitochondrial-encoded subunits show parallel elevation in the rates of evolution of nuclear-encoded subunits (Osada and Akashi 2012; Zhang and Broughton 2013; Sloan et al. 2014; Adrion et al. 2016; Havird et al. 2017). For example, studies in animals and yeast indicate that the scaled rates of evolution of mitochondrial ribosomal-protein sequences are *>* 10*×* those for cytoplasmic ribosomes (Pietromonaco et al. 1986; Barreto and Burton 2013; Barreto et al. 2018). Despite both sets of genes being encoded in the nuclear genome, they respectively assemble around mitochondrial- and nuclear-encoded ribosomal RNAs. Similar observations have been made for the complexes involved in organelle-harbored OXPHOS (oxidative phosphorylation) complexes, but notably, components of complexes that are fully encoded in the nuclear genome do not exhibit elevated rates of evolution (Havird et al. 2015; Weaver et al. 2022). Such coordinated patterns are expected if rate elevations in nuclear-encoded proteins are driven by the need to compensate for fixations of mildly deleterious alleles within a low-*N*, nonrecombining mitochondrion (which might also experience relaxed selection, i.e., reduced *s*).

Despite their high levels, *d_N_ /d_S_* for nuclear-encoded mitochondrial genes do not typically exceed the conventional criterion for positive selection (*d_N_ /d_S_* = 1.0). However, a few examples of *d_N_ /d_S_ >* 1.0 for nuclear-encoded subunits have been inferred to be associated with elevated organelle mutation rates (Osada and Akashi 2012; Rockenbach et al. 2016; Havird et al. 2017; Barreto et al. 2018). If these enhanced rates are indeed consequences of coevolutionary drive (as opposed to being results of other forms of positive selection), as noted above, the population-genetic environments of the two genomes must differ in a specific way with respect to *N* and *u*, with *Ns* being in the range of effective neutrality for the organelle subunits and in the range of efficient selection for the nuclear-encoded gene, and *u* being elevated in the organelle relative to the nuclear genome (Figures 3D and 4C). These observations are complemented by observations on plant-organelle genome evolution. For reasons that remain unexplained, in most land plants plastid genomes exhibit mutation rates that are 5 to 10*×* lower than those in coexisting nuclear genomes (Gaut et al. 1996), and the situation can be even more extreme in plant mitochondria. Notably, however, a few land-plant lineages have evolved dramatic increases in plastid mutation rates, and these exhibit the kinds of alterations in protein-sequence evolution noted above for metazoan mitochondria, including enhanced rates of amino-acid substitutions in nuclear-encoded subunits of plastid molecular complexes (Sloan et al. 2014; Zhang et al. 2015; Rockenbach et al. 2016; Weng et al. 2016).

As mentioned above, however, the unusual situation for land plants in which organelle genes (O) commonly have lower *N* and *u* relative to their nuclear-encoded (N) counterparts yields a substantially different prediction from that outlined in Figure 3D. In the generic land-plant case, the organellar site, with lower *N* and *u*, is expected to consistently have the higher *d_N_ /d_S_,* and neither site ever has *d_N_ /d_S_ >* 1 (Figure 5). For *N_N_ s <* 1, both sites evolve in an effectively neutral fashion, whereas for the intermediate regime (*N_N_ s >* 1 but *N_O_s <* 1), there is a shoulder in which the slowly mutating organellar site evolves in an effectively neutral manner, with the nuclear-encoded site compensating for the former but at a correspondingly lower *d_N_ /d_S_* owing to its higher mutation rate. In the illustrated example, the nuclear gene has a 100-fold higher mutation rate, resulting in a 100-fold reduction in *d_N_ /d_S_* in this intermediate region. Only after the effective population sizes at each site exceed 1*/s* does *d_N_ /d_S_* uniformly decline at both sites with increasing *N*.

**Figure 5.**
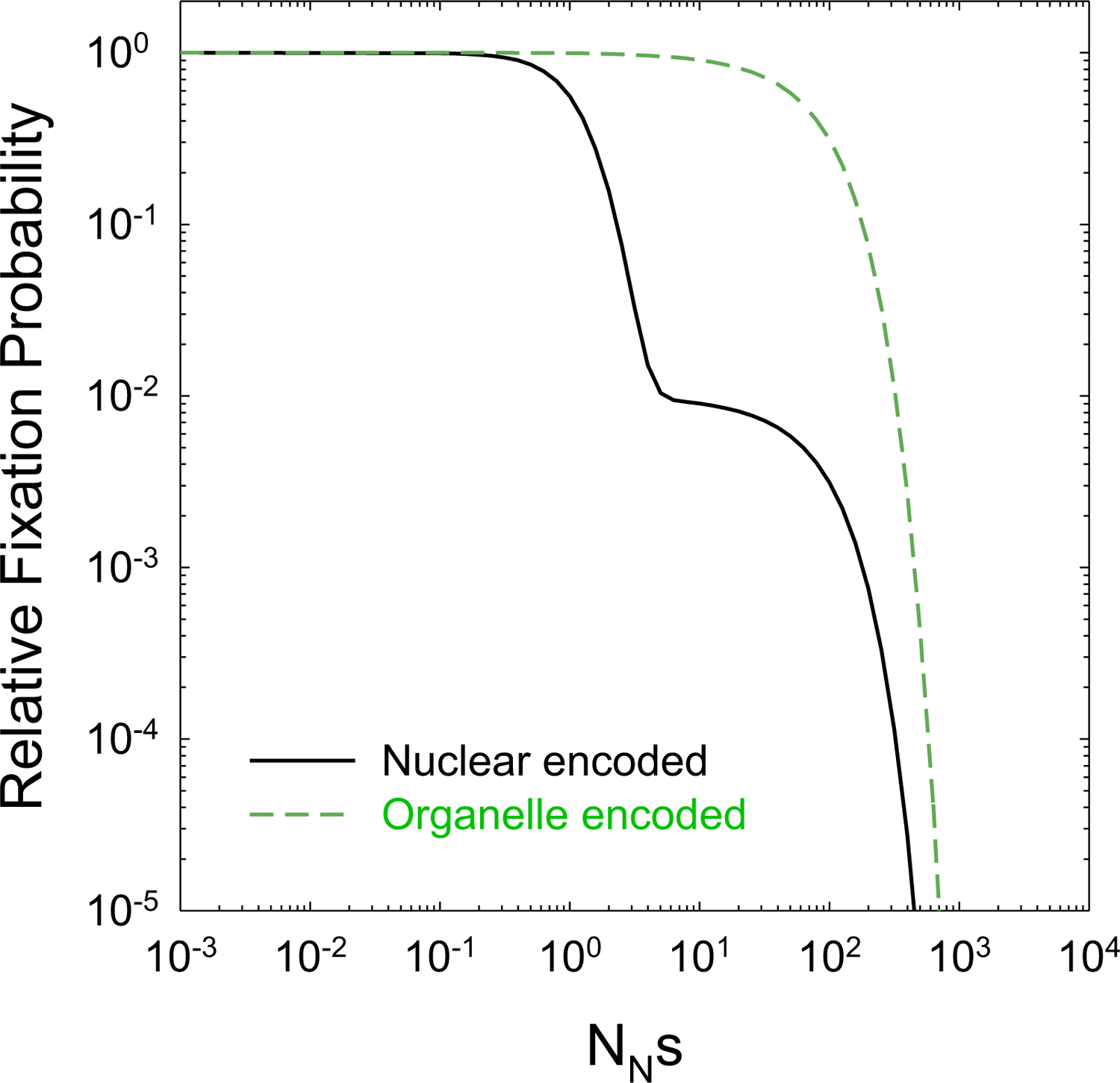
The behaviour of scaled fixation rates with unlinked loci when the mutation rate and effective population size is 100x lower at one site (green dashed line; *u* = 10^-9^) than at the other (black line).

These results suggest that comparative analyses involving interacting organelle/nuclear-encoded partners in species with different mutation-rate profiles should provide a useful basis for testing the theory outlined above. Indeed, many of the apparent inconsistencies between observations on compensatory mutations / molecular coevolution and verbal theoretical expectations (e.g., Piccinini et al. 2021) will remain unresolved until the population-genetic environments of the participating loci are taken into consideration. The following predictions are testable with sequence data from appropriately chosen phylogenetic lineages:

1. With one exception mentioned below, there should generally be a decline in *d_N_ /d_S_* with increasing *N*, i.e., coevolutionary constraints usually slow down the rate of evolution in increasingly large-*N* settings. This, of course, is the usual expectation for nucleotide sites under purifying selection, but in this case, there is an expected phylogenetic correlation between pair members, with both participating sites expected to have identical long-term scaled rates of evolution.
2. For completely linked sites, the behavior of the system should be independent of the mutation rate, unless *Ns >* 5, in which case however, *d_N_ /d_S_* is likely to be nearly indistinguishable statistically from 0.0 unless *u >* 10*^−^*^7^ or so.
3. When disparities in *N* exist between two coevolving sites, the site with the lower *N* is expected to dictate the pace of evolution, with *d_N_ /d_S_ ∼* 1.0 until *Ns >* 1 for the low *N* site.
4. When both *N* and the mutation rate (*u*) differ between coevolving sites, the behavior of the system qualitatively depends on the coupling of *N* and *u*. When the latter are negatively associated (e.g., low *N* with high *u*), the high-*N* /low-*u* site has the potential to exhibit *d_N_ /d_S_ >* 1, the only situation in which intraspecies molecular coevolution accelerates the rate of evolution beyond the neutral expectation. With the opposite coupling, the high-*N* /high-*u* site is expected to show substantially reduced *d_N_ /d_S_* relative to the alternative site, which never has *d_N_ /d_S_ >* 1.0.

Finally, it should be noted that the theory presented here is focused on situations in which molecular coevolution is governed by relatively constant internal cellular constraints, as might be expected for genes involved in transcription, translation, central metabolism, and intracellular signaling. Coevolution driven by external environmental forces that are themselves evolvable, e.g., mutualisms such as plant-pollinator and endosymbiotic bacterial-insect interactions (Futuyma and Slatkin 1983; Thompson 2005; Hembry et al. 2014; Week and Nuismer 2019), will require modifications of the model to include factors such as generation-time differences, nonobligatory interactions, and asymmetric fitness effects in the two interacting species (with the need to implement up to six selection coefficients).

However, given the restriction of the coevolution of unlinked genes to the sequential-model regime, the paths to extending the theory to many forms of interspecies molecular coevolution seems straight-forward.

## Materials and Methods

The work contained herein is largely based on analytical approximations of stochastic population-genetic processes. To determine the accuracy of such formulations, the underlying processes were simulated with a discrete-generation-time model incorporating consecutive generations of reversible mutation, recombination, selection, and random genetic drift operating on a two-locus biallelic system. The analyses were performed over the full range of biologically plausible parameter space, using self-written computer code, which compiles the long-term steady-state probability distribution of the four haplotypes as well as the rates of transition among them. The C++ program, TwoSites.cpp, is available at https://github.com/LynchLab/Molecular-Coevolution.

## Acknowledgments

This work was supported by National Institutes of Health grant R35-GM122566-01, US Department of Army MURI award W911NF-14-1-0411, National Science Foundation grants MCB-1518060 and DBI-2119963, and grant 735927 from the Moore and Simons Foundations. I am grateful to reviewers for helpful comments.

## SUPPLEMENTAL MATERIAL

Analytical results are provided here for some special cases, all of which derive from the general approaches described in the main text.

### Complete linkage; intermediate states with different deleterious effects; end states with equivalent high fitness

For the case of completely linkage, it is assumed that the sites have identical cellular and population-genetic environments so that population sizes and mutation rates are equal *N* = *N_A_* = *N_B_* and *u* = *u_A_* = *u_B_.* In this case, *s*_2_ */*= *s*_3_ *>* 0 and *s*_4_ = 0. Using Equations 1a-d and 3a,b, both end states have equal steady-state probabilities under the sequential model,

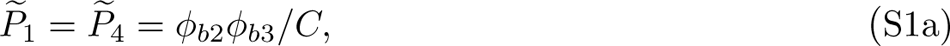

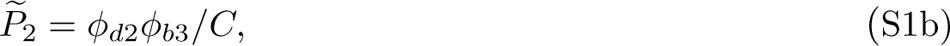

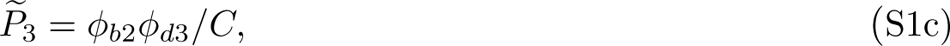

with the *φ* terms denoting fixation probabilities (b for beneficial, d for deleterious, and the number denoting the selection coefficient), and *C* being equal to the sum of the four numerators.

Both sites have equal rates evolution relative to the neutral expectation. From Equations 5a,b, for the sequential domain,

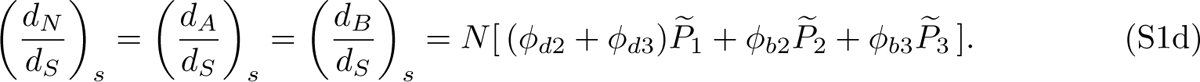

Stochastic tunneling operates through the low-fitness intermediate states **aB** and **Ab**, which in large populations are maintained at selection-mutation balance frequencies *u/s*_2_ and *u/s*_3_, resepctively. With *N* individuals and equal end-state fitnesses, the rate of fixation of tunneling mutations is equal to the product of these frequencies and *Nu ·* (1*/N*), leading to

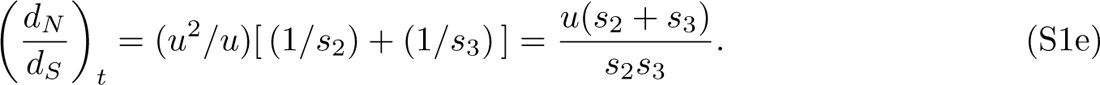

Allowing for the fact that effective tunneling requires large *Ns*, more generally,

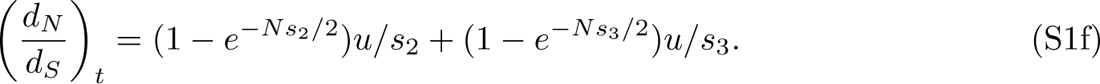

The total relative rate of evolution is then equal to the sum of Equations S1d and S1f.

### Complete linkage; intermediate states with equivalent deleterious effects; end states with elevated but different fitnesses

In this case, the selective disadvantages of the two intermediate states are *s* = *s*_2_ = *s*_3_ *>* 0, whereas one end state has advantage *s*_4_,. Given their linkage in the same genome, both sites are again assumed to have identical mutation rates and effective population sizes. Using Equations 1a-d and 3a,b, the steady-state probabilities under the sequential model are,

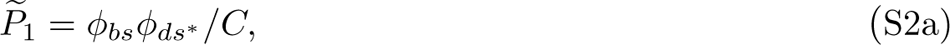

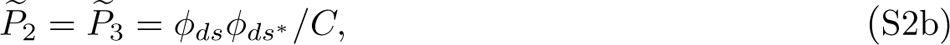

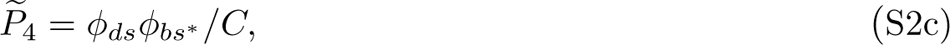

where *s^∗^* = *s* + *s*_4_, with the *φ* terms again denoting fixation probabilities (b for beneficial, d for deleterious), and *C* being equal to the sum of the four numerators.

From Equations 5a,b, for the sequential domain, both sites are again found to have equal rates of evolution relative to the neutral expectation.

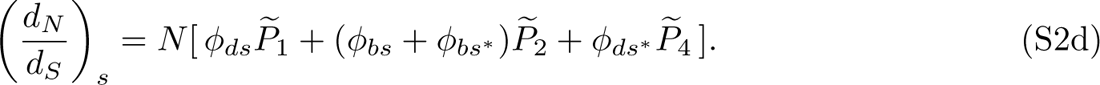

The rate of stochastic tunneling can be approximated by

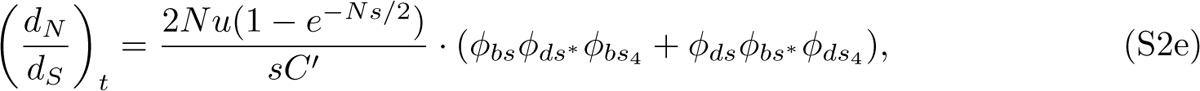

where *C^t^* is the sum of the numerators in Equations S2a and S2c. As *N → ∞,* the tunneling rate approaches (2*Nu/s*)*φ_ds_*_4_, which asymptotically approaches zero, owing to the fact that the population is permanently retained in the near-pure beneficial end state.

The total relative rate of evolution is equal to the sum of Equations S2d and S2e.

### Free recombination; symmetrical fitnesses (*s*_2_ = *s*_3_ = *s* and *s*_4_ = 0); arbitrary population sizes (*N_A_* and *N_B_*) and mutation rates (*u_A_* and *u_B_*)

The system of Equations 1a-b reduce to Equations 4a,b in the main text, and the rates of nucleotide substitution scaled to the neutral expectation are given by Equations 6a,b in the main text. The latter then expand to

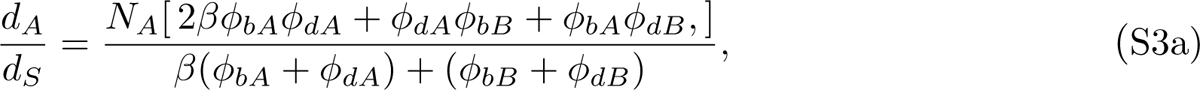

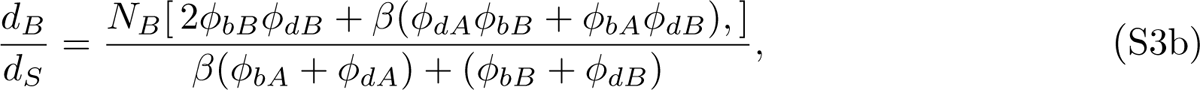

where *β* = (*N_A_u_A_*)*/*(*N_B_u_B_*), and the fixation probabilities are defined by Equations 3a,b using the appropriate population size. For *N_A_s* and *N_B_s »* 1, these expressions reduce to Equations 10a,b in the main text.

For the special case in which mutation rates differ, but population sizes are constant, then *φ_b_* = *φ_bA_*= *φ_bB_*, *φ_d_* = *φ_dA_*= *φ_dB_*, and Equations S3a,b reduce to Equation 7 in the main text.

### Free recombination; asymmetrical fitnesses (*s*_2_ */*= *s*_3_, but *s*_4_ = 0); constant population sizes (*N_A_* = *N_B_* = *N*); and arbitrary mutation rates (*u_A_, u_B_*)

The scaled evolutionary rates of both sites are identical and independent of the mutation rates,

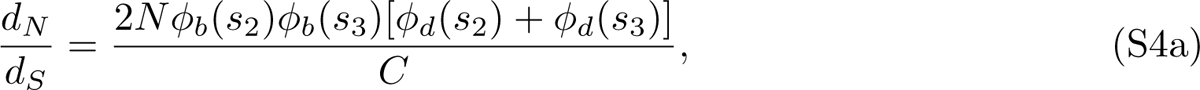

where

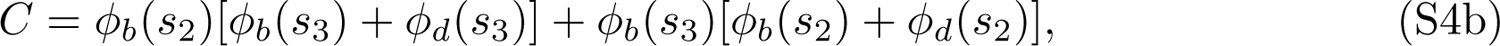

and the fixation probabilities are defined by Equations 3a,b using fixed *N* and substituting the appropriate selection coefficient.

---

**Supplemental Figure S1.**
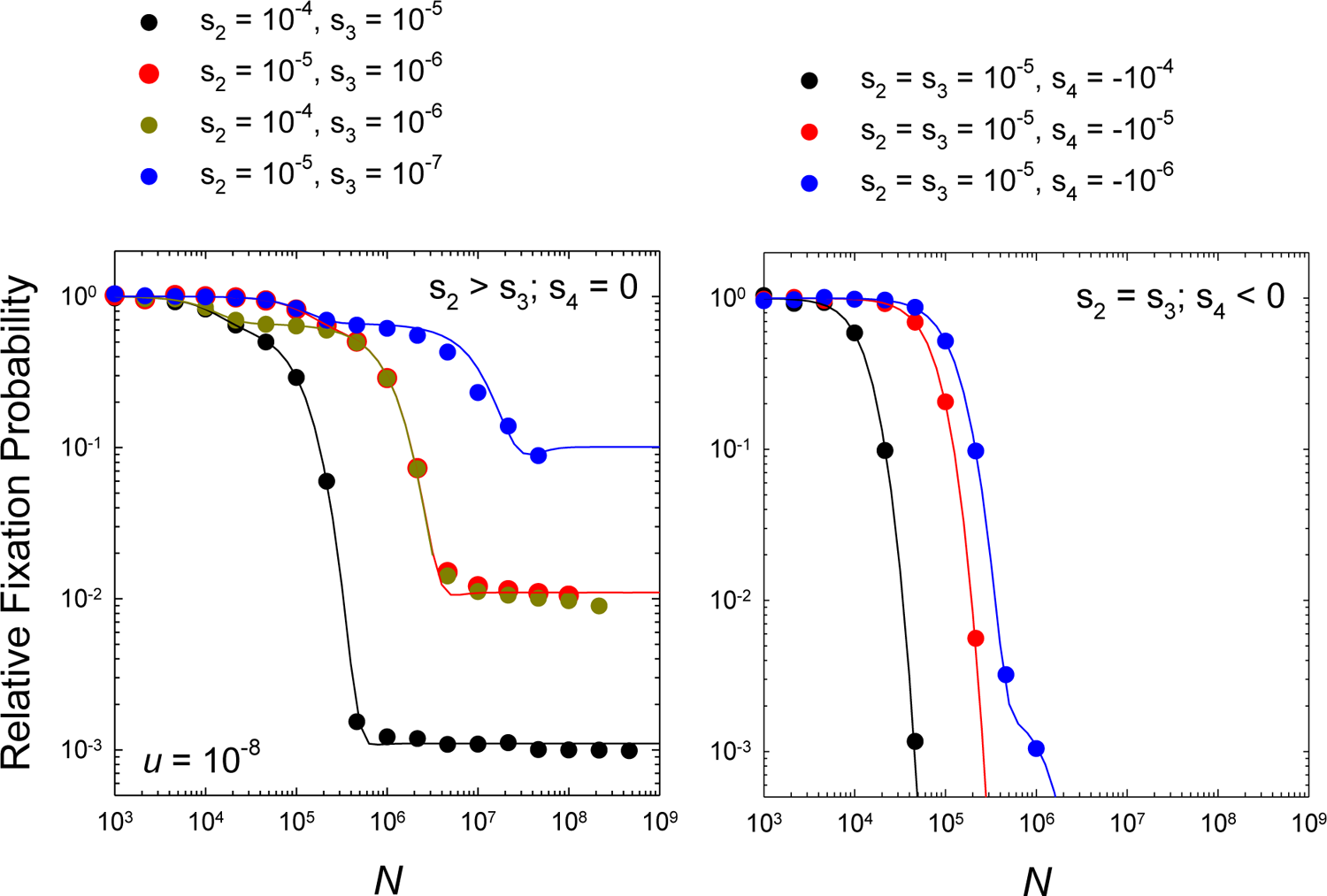
Rates of substitution at the two sites scaled to the neutral expectation as a function of *N*, for the case of complete linkage with both sites experiencing equal mutation rates and effective population sizes. The data points are computer-simulation results, obtained over a range of *N* for various combinations of *s_2_, s_3_*, and *s_4_*. The solid lines denote the predictions from theory outlined in the Supplemental Text, Equations S1d and S1f on the left, and Equations S2d and S2e on the right.

**Supplemental Figure S2.**
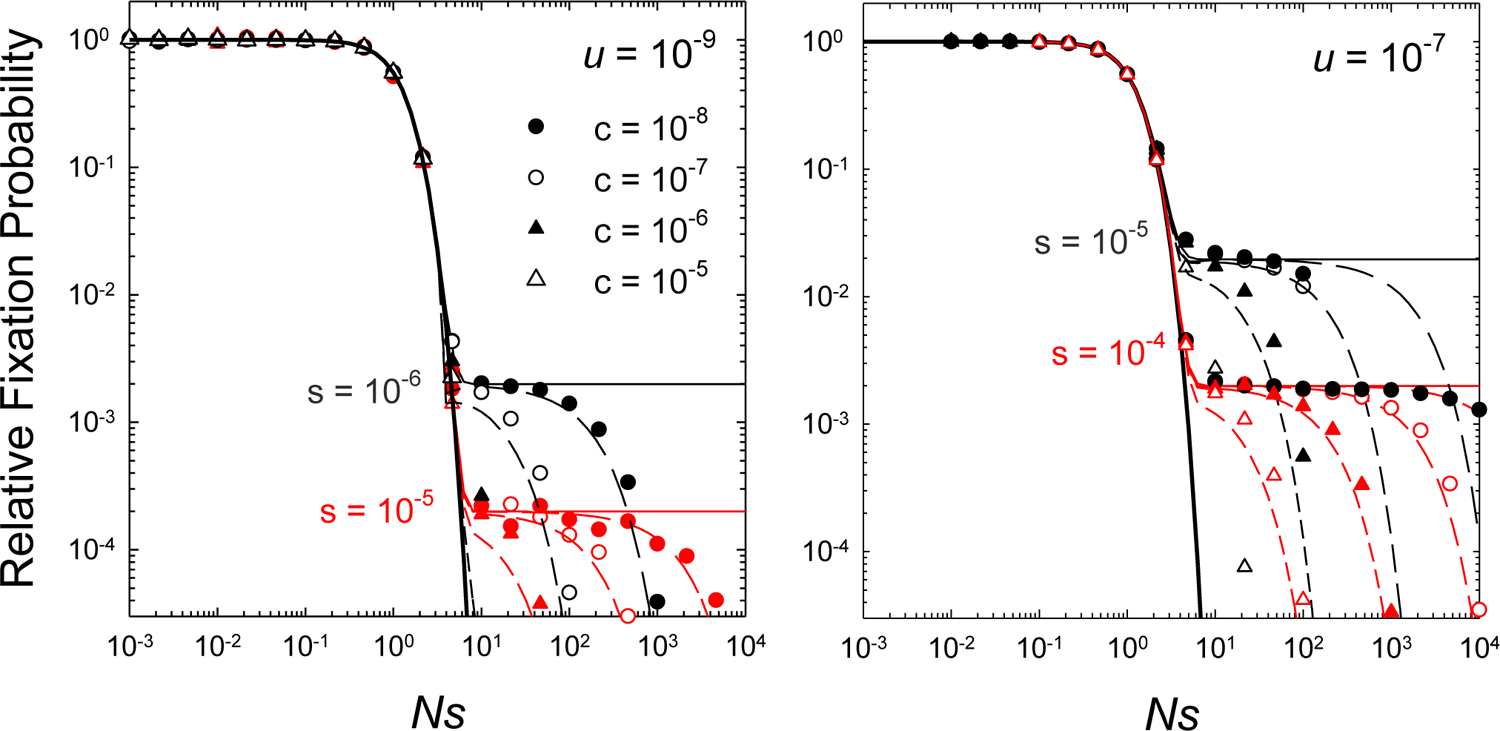
Influence of the recombination rate on the scaled rate of evolution. The solid lines are the theoretical expectations in the absence of recombination (as shown in Figure 2), whereas the curved lines are the theoretical expectations (described in the text) for intermediate levels of recombination (symbol shapes denoted in the inset, for two values of *s*). The data points are from computer simulations with fixations recorded as transitions of one haplotype to another using a threshold frequency of 0.999.

